# Unc93b1 recruits Syntenin-1 to dampen TLR7 signaling and prevent autoimmunity

**DOI:** 10.1101/409862

**Authors:** Olivia Majer, Bo Liu, Nevan Krogan, Gregory M. Barton

## Abstract

Recognition of nucleic acids enables detection of diverse pathogens by a limited number of innate immune receptors but also exposes the host to potential autoimmunity. At least two members of the Toll-like receptor (TLR) family, TLR7 and TLR9, can recognize self RNA or DNA, respectively. Despite the structural and functional similarities between these receptors, their contribution to autoimmune diseases such as SLE can be quite different. However, mechanisms of negative regulation that differentiate between TLR7 and TLR9 have not been described. Here we report a new function for the TLR trafficking chaperone Unc93b1 that specifically limits TLR7 signaling and prevents TLR7-dependent autoimmunity. Unc93b1 is known to traffic TLRs from the endoplasmic reticulum to endosomes, but this new regulatory function does not affect TLR7 localization. Instead, Unc93b1 recruits Syntenin-1, which inhibits TLR7, but not TLR9, signaling. Syntenin-1 binding requires phosphorylation of two serine residues on Unc93b1, providing a mechanism for dynamic regulation of the activation threshold of TLR7. Disruption of the Unc93b1/Syntenin-1 interaction in mice results in TLR7-dependent autoimmunity. Thus, Unc93b1 not only enables proper trafficking of nucleic acid sensing TLRs but also sets the activation threshold of these potentially self-reactive receptors.

## Main

Localization of nucleic acid-sensing TLRs to endosomes favors recognition of microbial versus self-derived ligands^1-3^. Several mechanisms cooperate to reinforce the compartmentalized activation of these receptors. First, the extracellular domains of nucleic acid-sensing TLRs must undergo proteolytic processing by endosomal proteases to generate a functionally competent receptor^4-6^. Second, nucleases in both the extracellular space and in endosomes degrade self-nucleic acids, reducing the likelihood that sufficient levels of ligand accumulate in endosomes and activate TLRs^7-11^. Finally, the amount of receptor in endosomes is carefully regulated via expression and sub-cellular trafficking^12-16^.

Altering any one of these general mechanisms can lead to autoimmunity, indicating that endosomal TLR signaling is carefully balanced to enable discrimination between self and foreign nucleic acids^3^. In many cases, though, the molecular pathways that control the expression, trafficking, and signaling of these potentially self-reactive TLRs have not been defined despite the clear relevance of such pathways to human disease. For example, TLR7 and TLR9 have opposing effects in mouse models of systemic lupus erythematosus (SLE); disease is exacerbated in TLR9-deficient mice but attenuated in TLR7-deficient mice^17,18^. Furthermore, *Tlr7* gene duplication can induce SLE in mice^12,15,16^, while overexpression of TLR9 causes little or no disease^19^. These observations suggest unique mechanisms of TLR7 and TLR9 regulation; however, mechanisms that distinguish endosomal TLR signaling are unknown.

Here we describe a new pathway mediated by the TLR trafficking chaperone Unc93b1 that differentially regulates the activation threshold of endosomal TLRs. Unc93b1 binds a subset of TLRs (TLR3, TLR5, TLR7, TLR8, TLR9, TLR11, TLR12, and TLR13) in the endoplasmic reticulum (ER) and facilitates their trafficking to endosomes^20-22^. Mice and humans lacking Unc93b1 have defective TLR function and exhibit increased susceptibility to certain viral infections^23^. Mutations in Unc93b1 can also enhance the responses of specific TLRs, leading to autoimmunity^13,14^. These examples of Unc93b1-mediated modulation of TLR function have been attributed to alterations of TLR trafficking^13,14^. Here we demonstrate a new mechanism by which Unc93b1 directly influences TLR7 signaling. We show that Unc93b1 recruits Syntenin-1 which dampens TLR7 signaling and limits responses to self RNA. Our work defines new functions of Unc93b1 and provides a mechanism for differential regulation of signaling by TLR7 and TLR9.

### Identification of a region within Unc93b1 that specifically regulates TLR7 responses

Unc93b1 is a twelve-pass transmembrane protein required for the function of all endosomal TLRs as well as TLR5^22,24,25^, yet mechanistic details regarding Unc93b1 function remain scarce. To identify regions of the protein involved in TLR regulation we performed a triple alanine mutagenesis screen of Unc93b1 using a library of 204 mutants covering the N-and C-terminal tail regions as well as all the loops. Each mutant was stably expressed in a RAW macrophage cell line in which both endogenous *Unc93b1* alleles were disrupted by Cas9 genome editing. As expected, deletion of endogenous Unc93b1 led to lack of responses to nucleic acids (Fig. S1a) and failure of TLR7 to traffic to endosomes (Fig. S1b). To evaluate TLR function in cells expressing each mutant, we stimulated each line with ligands for TLR3, TLR7, and TLR9 (Unc93b1-dependent TLRs) and TLR4 (an Unc93b1-independent TLR) and measured TNFα production. Complementing with Unc93b1^WT^ rescued signaling of TLR3, TLR7, and TLR9, while Unc93b1^H412R^, a previously described null allele^25^, did not (Fig. 1a). These analyses identified several Unc93b1 mutant alleles (PRQ(524-526)/AAA, PKP(530-532)/AAA, DNS(545-547)/AAA and DES(548-550)/AAA) that enhanced TLR7 responses relative to cells expressing wildtype Unc93b1, without affecting TLR3 or TLR9 responses (Figs. 1a,b). These mutations were all located within a 33 aa region in the Unc93b1 C-terminal tail (residues 521 to 553) (Fig. 1d), suggesting that the phenotypes associated with these mutants may be linked through a common mechanism. TLR7 responses were enhanced in response to the synthetic ligand R848 as well as to single-stranded RNA, especially when cells were stimulated with low concentrations of R848 (Fig. 1c). This enhanced signaling was not due to differences in the expression or stability of the Unc93b1 mutants, as protein levels were similar among the RAW macrophage lines (Fig. S1c).

**Fig. 1.**
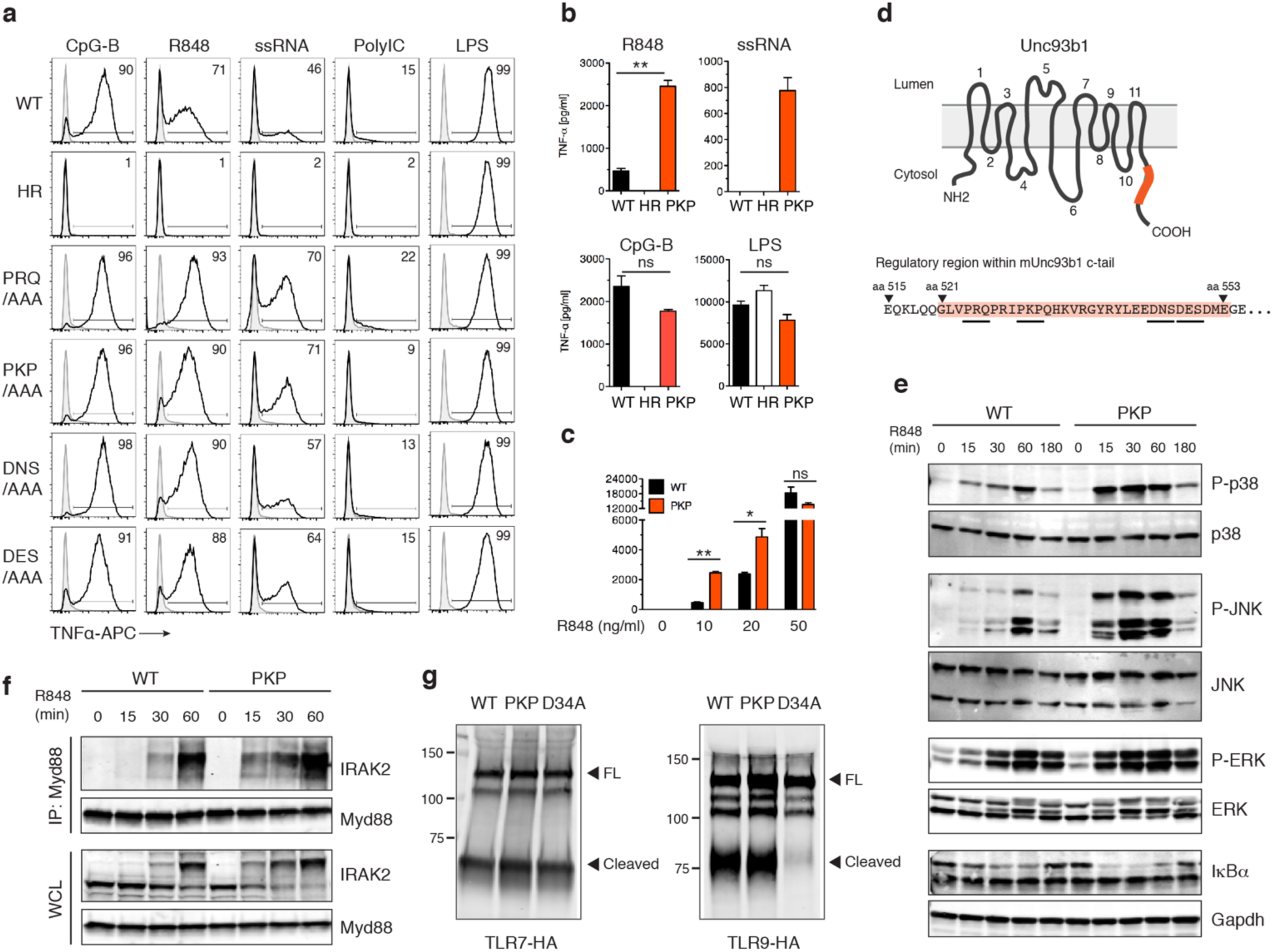
A C-terminal region in Unc93b1 regulates TLR7 responses. (**a**) Unc93b1 C-terminal mutants show enhanced TLR7 responses. Representative flow cytometry analysis showing percent TNFα positive cells, measured by intracellular cytokine staining, of Unc93b1-deficient RAW macrophages retrovirally transduced to express the indicated Unc93b1 alleles after stimulation with CpG-B (100 nM) for TLR9, R848 (10 ng/ml) and ssRNA40 (2.5 µg/ml) for TLR7, PolyIC (20 µg/ml) for TLR3, or LPS (10 ng/ml) for TLR4. (**b,c**) TNFα production, measured by ELISA, from the indicated RAW macrophage lines after stimulation for 8h with R848 (10 ng/ml), ssRNA40 (1 µg/ml), CpG-B (25 nM), LPS (50 ng/ml), or increasing concentrations of R848 (n=3; representative of three independent repeats). (**d**) Domain structure of Unc93b1 with the C-terminal regulatory region indicated in orange. (**e**) Unc93b1^PKP^-expressing macrophages show enhanced TLR7 signaling. Immunoblot of P-p38, P-JNK, P-ERK, and IκBα of RAW macrophages stimulated with R848 (50 ng/ml) for indicated times. Representative of two independent experiments. (**f**) Enhanced Myddosome assembly in Unc93b1^PKP^ macrophages. Immunoprecipitation of MyD88 from RAW macrophage lines expressing the indicated Unc93b1 alleles and stimulated with R848 (500 ng/ml), for indicated times followed by immunoblot for IRAK2. Input levels of Myd88 and IRAK2 in whole cell lysates (WCL) are also shown. (**g**) TLR7 and TLR9 trafficking are normal in Unc93b1^PKP^ but not in Unc93b1^D^34^A^ RAW lines. Immunoblot of TLR7 and TLR9 from lysates of indicated RAW macrophage lines. All data are mean ± SD; *p < 0.05, **p < 0.01, ***p < 0.001 by unpaired Student’s t-test. The data are representative of at least three independent experiments, unless otherwise noted.

To investigate how mutations in this region of Unc93b1 affect TLR7 signaling, we examined MAPK and NFκB activation in Unc93b1^PKP/AAA^ (hereafter referred to as Unc93b1^PKP^) expressing cells upon stimulation with R848. Activation of all three MAPKs (p38, JNK and ERK) was stronger and more rapid in Unc93b1^PKP^ cells, compared to Unc93b1^WT^ cells (Fig. 1e). Likewise, degradation of IκBα, a negative regulator of NFκB, occurred with faster kinetics in mutant cells (Fig. 1e). Assembly of the Myddosome complex, the most proximal signaling step downstream of TLR7 activation, also occurred more rapidly in Unc93b1^PKP^ expressing cells (Fig. 1f). Taken together, these results indicate that the C-terminal tail of Unc93b1 negatively regulates TLR7 signaling and presumably targets an early signaling event downstream of the receptor.

A previously described mutation (D34A) near the N-terminus of Unc93b1 that enhances TLR7 signaling has been attributed to increased TLR7 export from the ER at the expense of TLR9^14^, so we considered whether the same mechanism is responsible for the enhanced signaling in C-terminal mutants. However, trafficking of TLR7 and TLR9 appeared normal in Unc93b1^PKP^ expressing cells, as the cleaved form of both receptors, which represents the endosome-localized receptor, was present at levels comparable to Unc93b1^WT^ expressing cells (Fig. 1g). In contrast, the cleaved form of TLR9 was absent and TLR9 signaling was defective in Unc93b1^D^34^A^-expressing cells, consistent with the block in TLR9 trafficking reported for this mutant^14^ (Fig. S2). These results suggested that the C-terminal region of Unc93b1 specifically regulates TLR7 responses through a novel mechanism.

### Unc93b1^PKP^ leads to enhanced TLR7 signaling without altering TLR7 trafficking or localization

We next investigated the mechanism(s) by which mutations in the Unc93b1 C-terminal tail lead to enhanced TLR7 responses. We considered the possibility that more TLR7 is exported from the ER to endosomes in Unc93b1^PKP^-expressing cells, similar to the mechanism proposed for Unc93b1^D^34^A^-expressing cells^14^, but multiple observations argued against this model. As noted earlier, we did not observe any increase in the steady-state level of cleaved TLR7, which represents the endosomal pool of the receptor, in Unc93b1^PKP^-expressing RAW cells (Fig. 1g). Also, pulse/chase analysis of TLR7 showed that ectodomain cleavage of TLR7 occurred with similar kinetics in Unc93b1^WT^-and Unc93b1^PKP^-expressing cells (Fig. 2a), suggesting that TLR7 trafficking to endosomes is equivalent. Finally, TLR7 levels were equivalent in phagosomes isolated from Unc93b1^WT^-and Unc93b1^PKP^-expressing cells (Fig. 2b).

**Fig. 2.**
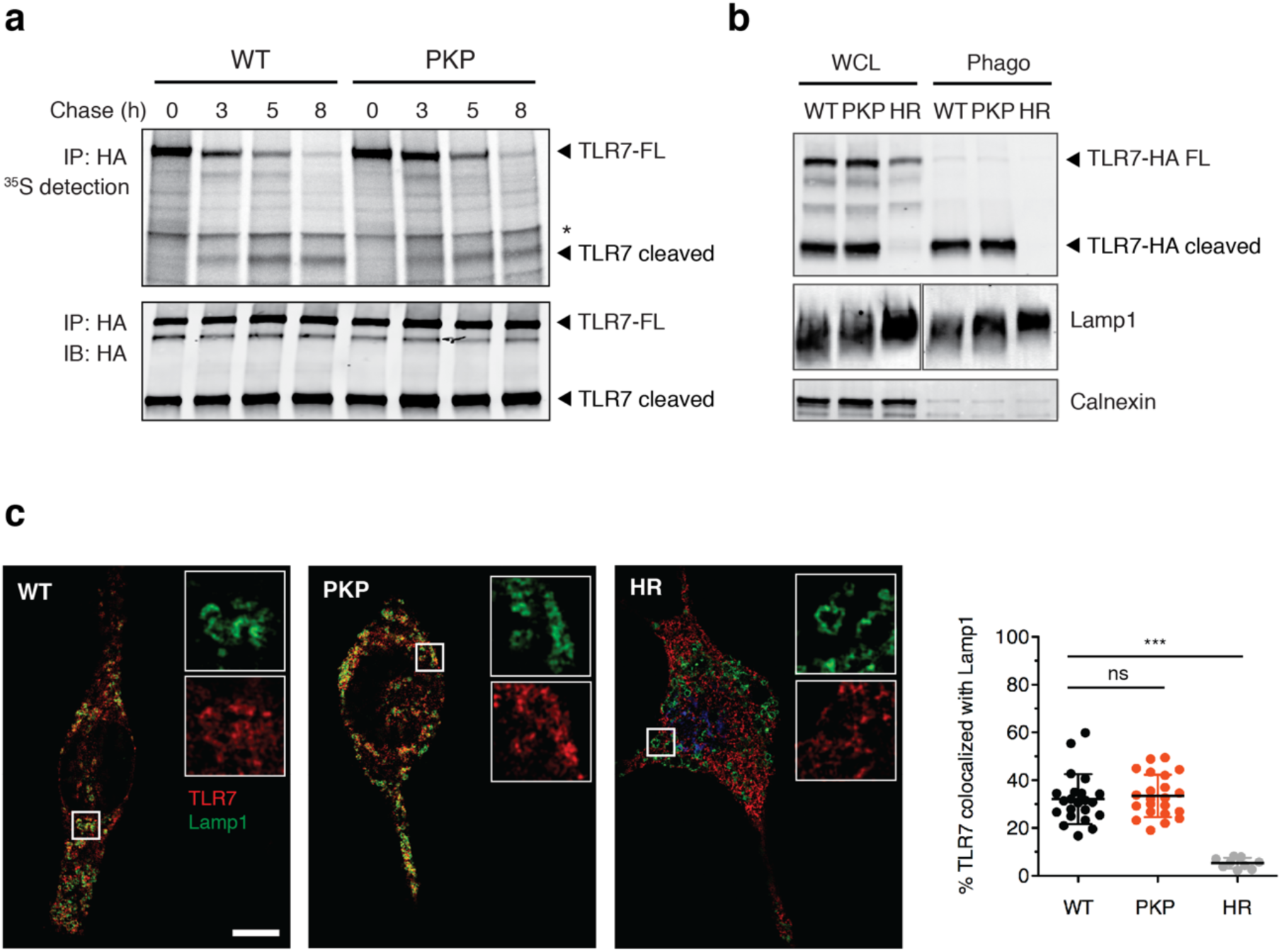
Unc93b1^PKP^ does not alter TLR7 trafficking or localization. (**a**) Unc93b1^PKP^ does not alter TLR7 export rates. Pulse–chase analysis of TLR7 in Unc93b1^WT^ and Unc93b1^PKP^-expressing RAW macrophages. Cell lysate was HA immunoprecipitated and subjected to radiolabeled screen and immunoblot. The full-length and cleaved forms of TLR7 are indicated. An asterisk denotes a nonspecific band. Representative of two independent experiments. (**b**) Unc93b1^PKP^ does not affect TLR7 trafficking to endosomes. Levels of TLR7, Lamp1, and Calnexin in whole cell lysates (WCL) or lysates of purified phagosomes from the indicated RAW macrophage lines were measured by immunoblot. Representative of three independent experiments. (**c**) Colocalization of TLR7 and Lamp1 in RAW macrophages expressing the indicated Unc93b1 alleles on a *Myd88*^*-/-*^ background using superresolution structured illumination microscopy. Shown are representative Unc93b1^WT^, Unc93b1^HR^ and Unc93b1^PKP^ cells: TLR7 (red) and Lamp1 (green). Boxed areas are magnified. The plot shows quantification of the percentage of total TLR7 within Lamp1^+^ endosomes with each dot representing an individual cell (pooled together from two independent experiments). Scale bars: 10 µm. All data are mean ± SD; *p < 0.05, **p < 0.01, ***p < 0.001 by unpaired Student’s t-test.

We also examined TLR7 and Unc93b1 localization directly using immunofluorescence microscopy. To avoid any aberrant results due to activation of TLR7 in cells ectopically expressing TLR7 and Unc93b1^PKP^ (e.g., from RNA released from dead cells in the well), we analyzed localization in cells lacking Myd88. The extent of colocalization between TLR7 and the late endosomal marker Lamp1 was similar in cells expressing Unc93b1^WT^ and Unc93b1^PKP^ but much reduced in cells expressing the null allele Unc93b1^H412R^ (Fig. 2c). There was also no apparent change in colocalization of Lamp1 with Unc93b1^PKP^ itself (Fig. S3). Altogether, these results suggest that the enhanced TLR7 response in Unc93b1^PKP^ cells cannot be explained by alterations in TLR7 trafficking and/or localization.

### Syntenin-1 binds to the Unc93b1 C-terminal tail and inhibits TLR7 signaling

Our findings thus far suggested that the Unc93b1 C-terminal tail might regulate TLR7 signaling in endosomes through a mechanism unrelated to its function as a trafficking chaperone, either by interfering with TLR7 signaling directly or by association with proteins that interfere with signaling. To test the latter possibility, we sought to identify proteins that interact with Unc93b1^WT^ but not Unc93b1^PKP^. One challenging aspect of this approach is the relatively small fraction (<5%) of Unc93b1 in endosomes relative to the ER (Figs. S3 and S4a). To overcome this obstacle, we enriched for the endosomal pool of Unc93b1 by isolating phagosomes from RAW cells expressing Unc93b1^WT^ or Unc93b1^PKP^ and then purified Unc93b1 protein complexes via anti-FLAG antibodies (see scheme in Fig. S4b). This approach revealed an approximately 32kDa band present in Unc93b1^WT^ samples that was reduced in Unc93b1^PKP^ samples (Fig. 3a). Using tandem mass spectrometry, peptide sequences from the Unc93b1^WT^ enriched band identified Syntenin-1 (also known as syndecan binding protein, SDCBP), which we confirmed by immunoblot, using an anti-Syntenin-1 monoclonal antibody (Fig. 3b).

**Fig. 3.**
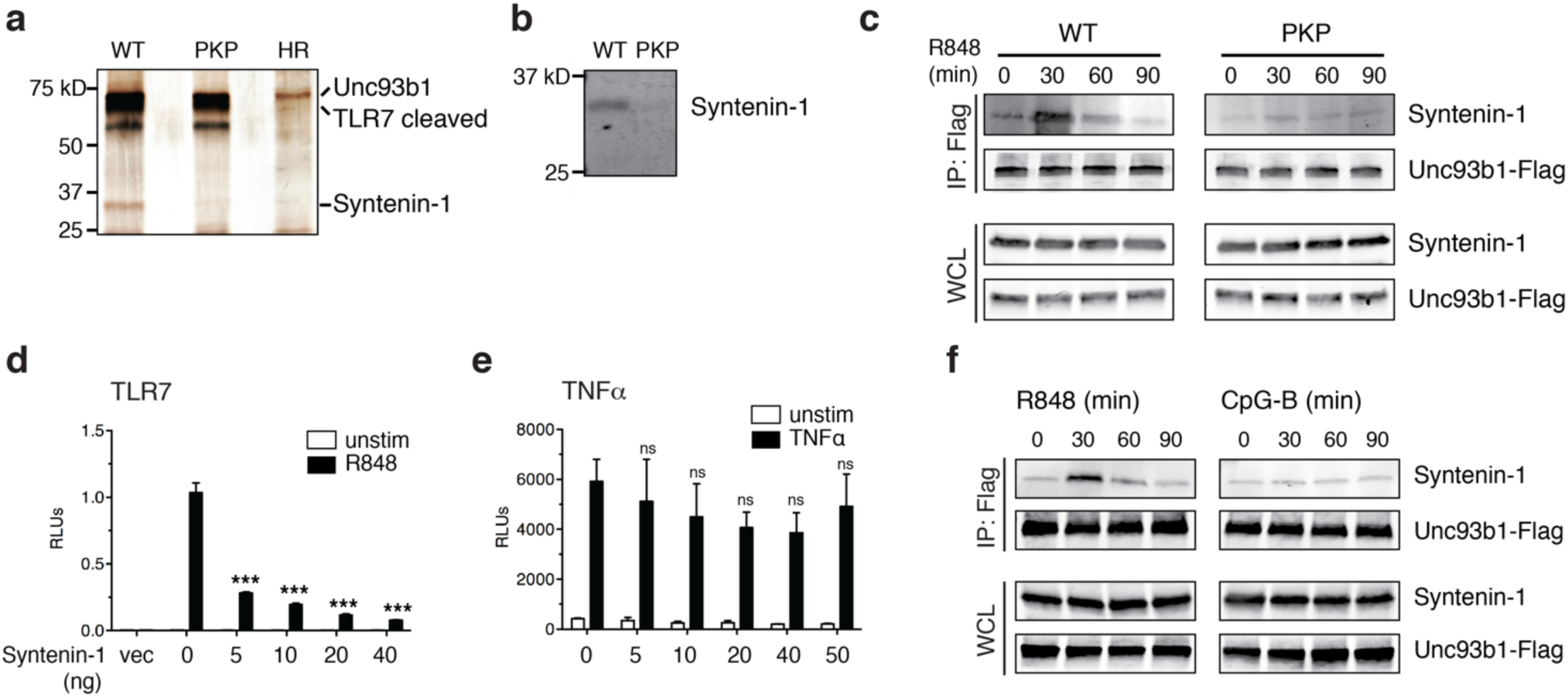
Syntenin-1 binds to the C-terminal tail of Unc93b1 and restricts TLR7 signaling. (**a**) Silver stained SDS-PAGE gel of purified Unc93b1-FLAG complexes in phagosomes enriched from RAW macrophages expressing the indicated Unc93b1 alleles. Note the 32kD protein (subsequently identified as Syntenin-1) that is less abundant in Unc93b1^PKP^ complexes. (**b**) Syntenin-1 binds Unc93b1^WT^. Purified Unc93b1-FLAG complexes described in (**a**) were immunoblotted for Syntenin-1. (**c**) Syntenin-1 is recruited to wildtype Unc93b1 but not Unc93b1^PKP^. Syntenin-1 binding to Unc93b1 was measured by FLAG immunoprecipitation followed by immunoblot for Syntenin-1 from RAW macrophage lines stimulated with R848 (0.5 µg/ml) for the indicated times. Levels of Syntenin-1 and Unc93b1-FLAG in cell lysates are also shown. (**d-e**) Syntenin-1 suppresses TLR7 signaling. (**d**) NFκB activation in HEK293T cells transiently expressing TLR7 and increasing amounts of Syntenin-1 was measured using a dual luciferase reporter assay. Cells were stimulated with R848 (50 ng/ml) for 16 h prior to harvest. On-way ANOVA results: F(4/10)=48.4, *p*<0.0001 (**e**) HEK293T cells transiently expressing Syntenin-1 and stimulated with TNFα (10 ng/ml); F(5/12)=1.263, *p*=ns. Data are normalized to Renilla expression and expressed as relative luciferase units (RLU) (n=3-4, representative of three independent experiments). (**f**) Syntenin-1 is selectively recruited to Unc93b1 upon TLR7 stimulation. Interaction between Syntenin-1 and Unc93b1 was measured as described in (c) from Unc93b1^WT^ RAW macrophages stimulated with R848 (0.5 µg/ml) or CpG-B (0.5 µM) for the indicated times. All data are mean ± SD and were analyzed with one-way ANOVA followed by a Tukey’s posttest (95% confidence interval): *p < 0.05, **p < 0.01, ***p < 0.001. The data are representative of at least three independent experiments.

Syntenin-1 is an adaptor protein that can influence trafficking of transmembrane proteins^26^ but has also been shown to regulate assembly of signaling complexes, including signaling downstream of TLRs^27^. We could detect Syntenin-1 associated with Unc93b1 in unstimulated RAW macrophages, but the interaction was rapidly and transiently increased after stimulation with TLR7 ligand (Fig. 3c). Consistent with our proteomic analyses, Syntenin-1 association with Unc93b1^PKP^ was reduced in unstimulated cells and did not increase after stimulation with TLR7 ligand (Fig. 3c).

Based on these results, we considered the possibility that Syntenin-1 negatively regulates TLR7 when bound to Unc93b1. A previous study has suggested that Syntenin-1 can inhibit TLR4 and IL-1 receptor (IL-1R) signaling by interfering with the interaction between IRAK-1 and TRAF6, but whether Syntenin-1 could inhibit TLR7 signaling was not examined^27^. To test this possibility, we measured the effect of Syntenin-1 overexpression on TLR7 activation in HEK293T cells, using an NF-κB luciferase reporter system. Increasing expression of Syntenin-1 substantially reduced TLR7 signaling (Fig. 3d), whereas activation of NF-κB by TNFα was not affected (Fig. 3e).

Since Unc93b1^PKP^ had no effect on TLR9 signaling, we asked whether Syntenin-1 recruitment was specific to TLR7. The increased association between Unc93b1 and Syntenin-1, as seen before (Fig. 3c), only occurred after TLR7 activation, as stimulation with TLR9 ligand did not change the levels of Syntenin-1 bound to Unc93b1 (Fig. 3f). Presumably, the selective inhibition of TLR7 signaling is based on the interaction between Syntenin-1 with Unc93b1, which brings Syntenin-1 in close proximity to TLR7. In an accompanying paper, we demonstrate that TLR9 is released from Unc93b1 within endosomes while TLR7 and Unc93b1 remain associated^28^. This mechanism likely explains why the disruption of Syntenin-1 inhibition with the Unc93b1^PKP^ mutation does not impact TLR9. Altogether, these results identify Syntenin-1 as an Unc93b1-binding protein that specifically inhibits TLR7 signaling.

### Syntenin-1 recruitment to Unc93b1 is regulated through phosphorylation of serine residues in the Unc93b1 C-terminal tail

The transiently increased Syntenin-1 binding to Unc93b1 upon TLR7 stimulation prompted us to investigate mechanisms by which this association could be dynamically regulated. Several global analyses of cellular phosphorylated proteins have identified potential phosphorylation of two serine residues (Ser547, Ser550) within the Unc93b1 C-terminal tail^29^. Two triple alanine mutations that encompass these serines (*Unc93b1*^*DNS/AAA*^ and *Unc93b1*^*DES/AAA*^) were identified in our original screen as leading to enhanced TLR7 responses (Fig. 1a), so we considered whether the disruption of phosphorylation at Ser547 and Ser550 underlies the defective regulation observed for these mutants. In fact, mutation of Ser547 (Unc93b1^S^547^A^), Ser550 (Unc93b1^S^550^A^), or both serines (Unc93b1^S^547^A/S^550^A^) to alanine was sufficient to enhance TLR7 responses to levels comparable to Unc93b1^PKP^ (Figs. 4a-c). To examine directly whether Ser547 and Ser550 of Unc93b1 are phosphorylated in cells, we generated polyclonal antisera specific for phosphorylated Ser547 and Ser550 within the Unc93b1 C-terminal tail (Fig. S5a). Using affinity-purified IgG from these sera, we confirmed that Unc93b1 is phosphorylated at these residues when expressed in RAW cells; mutation of either serine reduced detection by the phospho-specific Unc93b1 antibodies, while mutation of both serines completely abrogated detection (Fig. S5b). Based on these results, we conclude that at least some fraction of Unc93b1 is phosphorylated at Ser547 or Ser550 and that phosphorylation of these residues is necessary to limit TLR7 responses.

**Fig. 4.**
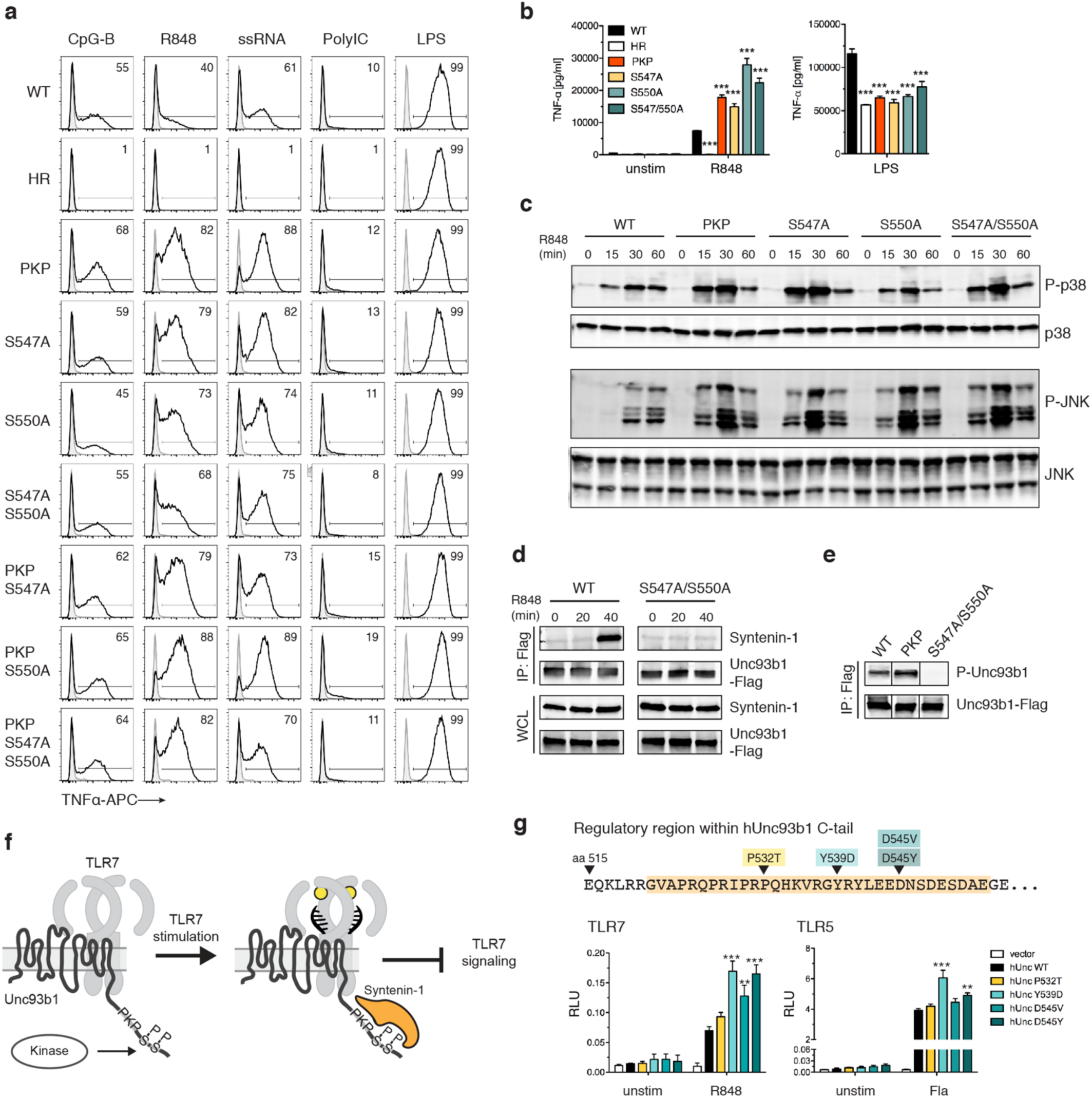
Serine phosphorylation in the C-terminal tail of Unc93b1 regulates Syntenin-1 recruitment. (**a**) Serine to alanine mutations in the C-terminal tail of Unc93b1 lead to enhanced TLR7 responses. Representative flow cytometry analysis showing percent TNFα positive cells, measured by intracellular cytokine staining, of Unc93b1-deficient RAW macrophages complemented with the indicated mutant alleles and stimulated with CpG-B (10 nM) for TLR9, R848 (10 ng/ml) and ssRNA40 (1 µg/ml) for TLR7, PolyIC (20 µg/ml) for TLR3, or LPS (10 ng/ml) for TLR4. (**b**) TNFα production, measured by ELISA, from the indicated RAW macrophage lines after stimulation for 8 h with R848 (20 ng/ml), or LPS (50 ng/ml) (n=3; representative of three independent experiments. One-way ANOVA results for R848 groups: F(5/12)=247.2, *p*<0.0001; LPS groups: F(5/12)=93.59, *p*<0.0001). (**c**) Levels of phospho-p38 and phospho-JNK, as measured by immunoblot, in lysates of the indicated RAW macrophage cells stimulated with R848 (50 ng/ml). (**d**) Phosphorylation of Ser547 and Ser550 is required for Syntenin-1 binding to Unc93b1. Syntenin-1 binding to Unc93b1^WT^ or Unc93b1^S^547^A/S^550^A^ in RAW macrophages stimulated with R848 (0.5 µg/ml) was measured by Unc93b1-FLAG immunoprecipitation followed by immunoblot for Syntenin-1. Levels of Syntenin-1 and Unc93b1-FLAG in cell lysates are also shown. (**e**) Unc93b1^PKP^ is phosphorylated. Unc93b1-FLAG was immunoprecipitated from lysates of the indicated RAW macrophage lines and phosphorylation of Ser547 and Ser550 was measured by immunoblot with phospho-specific antibodies. Each blot was performed on the same membrane but cropped to present relevant lanes. (**f**) A model of Syntenin-mediated TLR7 restriction. (**g**) Genetic variation in the human Unc93b1 C-terminal regulatory region increases TLR7 responses. NFκB activation in HEK293T cells transiently expressing TLR7 or TLR5 and the indicated human Unc93b1 alleles was measured using a dual luciferase reporter assay. Cells were stimulated with R848 (10 ng/ml) or ultrapure Flagellin (2 ng/ml) for 16 h prior to harvest. Data are normalized to Renilla expression and expressed as relative luciferase units (RLUs) (n=4, representative of three independent experiments. On-way ANOVA results for TLR7 groups: F(4/10)=20.03, *p*<0.0001; TLR5 groups: F(4/10)=28.69, *p*<0.0001). All data are mean ± SD; data were analyzed with one-way ANOVA followed by a Tukey’s posttest (95% confidence interval): *p < 0.05, **p < 0.01, ***p < 0.001. The data are representative of at least three independent experiments, unless otherwise noted.

Next, we sought to determine whether phosphorylation of Ser547 and Ser550 was inhibiting TLR7 signaling through the same mechanism that we uncovered for Unc93b1^PKP^. Combining the Unc93b1^PKP^ mutation with Unc93b1^S^547^A^, Unc93b1^S^550^A^, or Unc93b1^S^547^A/S^550^A^ did not further enhance TLR7 responses beyond that observed in Unc93b1^PKP^-expressing cells (Fig. 4a), suggesting that each mutation acts by disrupting the same mechanism. Indeed, Syntenin-1 recruitment to Unc93b1 after R848 stimulation was impaired in Unc93b1^S^547^A/S^550^A^-expressing cells (Fig. 4d), indicating that phosphorylation of Ser547 and Ser550 is required for binding of this negative regulator of TLR7 signaling.

In light of these results, the defect in Syntenin-1 recruitment associated with the Unc93b1^PKP^ mutation could either be due to disruption of residues required for direct interaction with Syntenin-1, or, alternatively, Unc93b1^PKP^ may fail to interact with the kinase(s) that phosphorylate(s) Ser547 and Ser550. To distinguish between these possibilities, we examined phosphorylation of Ser547 and Ser550 in Unc93b1^PKP^-expressing cells and observed that levels of phosphorylation were equivalent to the levels in Unc93b1^WT^ cells (Fig. 4e). These results support a model in which Syntenin-1 binding to Unc93b1 requires specific residues within the C-terminal tail as well as phosphorylation of Ser547 and Ser550 (Fig. 4f). While some Syntenin-1 is associated with Unc93b1 in unstimulated cells, the interaction is further increased upon TLR7 signaling. Thus, the mechanism we describe appears not only to influence the initial threshold of TLR7 activation but also operates as a negative feedback loop to shut down TLR7 signaling. The signals leading to this increased recruitment as well as the identities of the kinases and phosphatases regulating phosphorylation of Ser547 and Ser550 are important open questions for future work.

### SNPs in the Syntenin-1 binding region of human Unc93b1 can enhance TLR7 responses

The 33 aa region in the Unc93b1 C-terminal tail is highly conserved between mouse and human. To determine if modulation of Unc93b1/Syntenin-1 regulation of TLR7 signaling could be relevant in humans, we searched publicly available human genomic data for single nucleotide polymorphisms (SNPs) within the Unc93b1 C-terminal tail. Four very rare (Minor allele frequencies of 0.04% or lower, according to the 1000 Genomes project) coding variants within this region have been reported: P532T, Y539D, D545V, and D545Y (Fig. 4g). We tested for any functional consequences of these variants by expressing each in HEK293T cells together with TLR7. Three of the variants (Unc93b1^Y539D^, Unc93b1^D545V^, and Unc93b1^D545Y^) increased TLR7 responses relative to Unc93b1^WT^, although Unc93b1^Y539D^, and to a lesser extent Unc93b1^D545Y^, also increased TLR5 responses (Fig. 4g). These alleles are too rare to be linked to autoimmune disorders via genome-wide association studies, but the results suggest that modulation of the regulatory mechanism we describe here can influence TLR7 activation thresholds and consequently may impact the likelihood of certain autoimmune diseases.

### Mutation of the C-terminal regulatory region in Unc93b1 leads to TLR7-dependent autoimmunity

Finally, we sought to test the importance of Unc93b1/Syntenin-1 regulation of TLR7 for self versus non-self discrimination *in vivo.* We considered that analysis of Syntenin-1 deficient mice could be complicated for multiple reasons. First, Syntenin-1 has been implicated in the regulation of multiple transmembrane proteins, and Syntenin-1 deficient mice show pleiotropic effects on the immune system and the microbiota^30^. In addition, Syntenin-2, a highly homologous protein may compensate for Syntenin-1 deficiency. To sidestep these potential issues, we introduced the Unc93b1^PKP^ mutation into the germline of mice using Cas9 genome editing (Fig. S6a). This mutation disrupts interaction with Syntenin-1 but should leave other Syntenin-1 functions unaffected. We obtained an *Unc93b1*^*WT/PKP*^ founder, backcrossed this founder to C57BL/6J for 1 generation, and intercrossed *Unc93b1*^*WT/PKP*^ mice to generate *Unc93b1*^*WT/WT*^, *Unc93b1*^*WT/PKP*^, and *Unc93b1*^*PKP/PKP*^ offspring for analysis. *Unc93b1*^*PKP/PKP*^ mice were born below the expected Mendelian frequency and were severely runted (Fig. 5a). These mice exhibited hallmarks of systemic inflammation and autoimmunity, including increased frequencies of activated T cells, loss of marginal zone (MZ) B cells, increased frequencies of MHC^hi^ dendritic cells, and increased frequencies of inflammatory monocytes in secondary lymphoid organs (Fig. 5b). *Unc93b1*^*PKP/PKP*^ mice developed anti-nuclear antibodies (ANA) very early in life (Fig. 5c). *Unc93b1*^*WT/PKP*^ mice also showed signs of immune dysregulation but not to the same extent as *Unc93b1*^*PKP/PKP*^ mice (Figs. 5a-c).

**Fig. 5.**
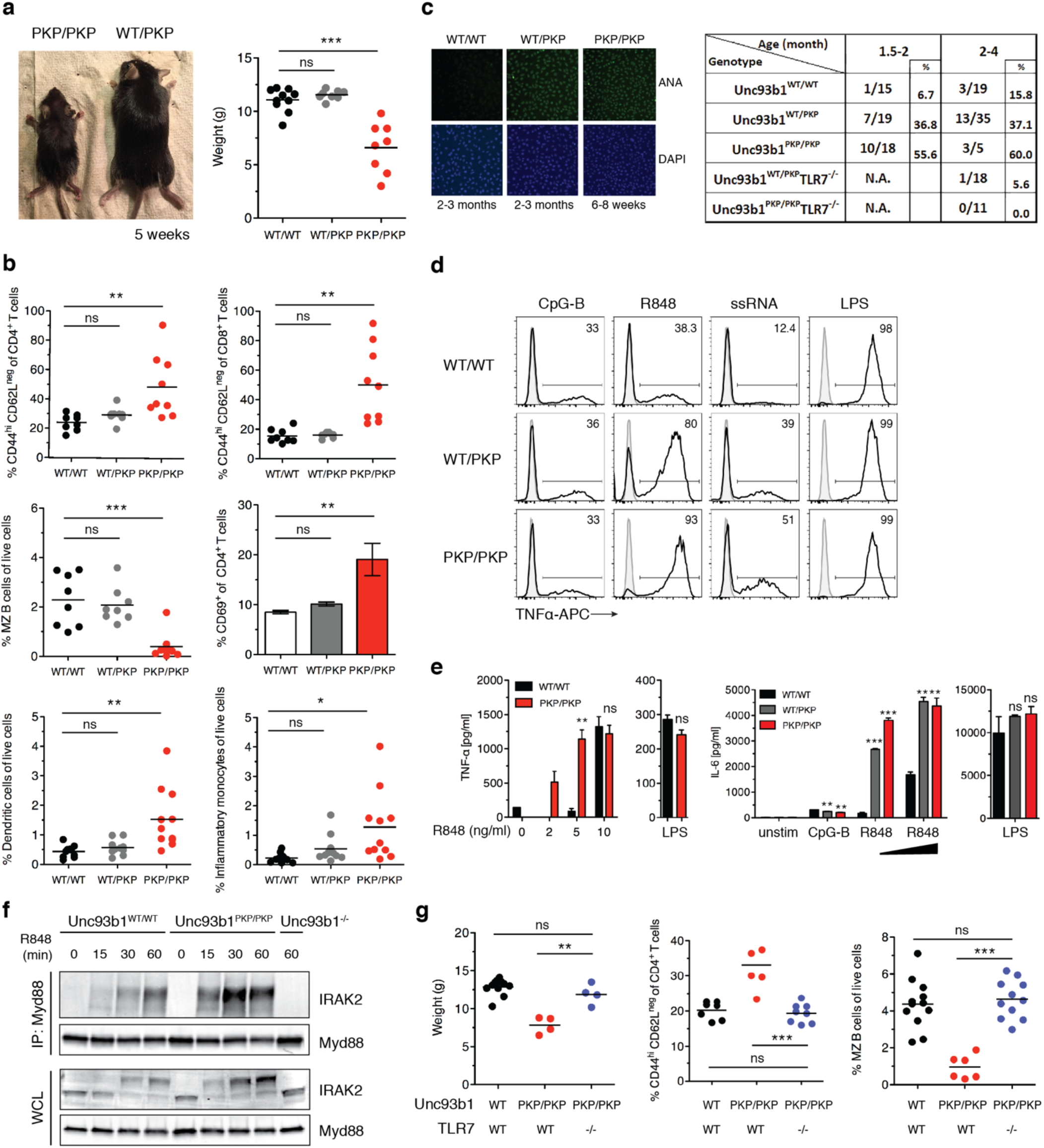
Unc93b1^PKP^ knock-in mice develop TLR7-driven systemic inflammation and autoimmunity. (**a**) Gross appearance and weights of *Unc93b1*^*PKP/PKP*^ mice compared to littermate controls. (**b**) *Unc93b1*^*PKP/PKP*^ mice exhibit systemic inflammation. Flow cytometry analysis of major immune cell subsets in *Unc93b1*^*WT/WT*^, *Unc93b1*^*PKP/WT*^, and *Unc93b1*^*PKP/PKP*^ mice at 6-8 weeks of age. Frequencies of CD44^high^ CD62L^neg^ of CD4^+^ T cells or of CD8^+^ T cells in spleen, marginal zone (MZ) B cells (CD21^+^CD23^neg^IgM^+^CD1d^+^) in spleen, CD69^+^ of CD4^+^ T cells in lymph nodes, dendritic cells (CD11b^+^ CD11c^+^ MHC-II^high^) in lymph nodes, and inflammatory monocytes (CD11b^+^Ly6c^+^Ly6G^neg^) in lymph nodes are shown. Data points were pooled from four independent experiments. (**c**) *Unc93b1*^*PKP/PKP*^ mice develop anti-nuclear antibodies (ANA) early in life. Representative staining (left) and tabulated results (right) of Hep-2 slides using sera (diluted 1:80) from the indicated mouse ages and genotypes. (**d**) Cells from *Unc93b1*^*PKP/WT*^ and *Unc93b1*^*PKP/PKP*^ mice show enhanced TLR7 responses. Representative flow cytometry analysis showing percent TNFα positive cells, measured by intracellular cytokine staining, of bone marrow-derived dendritic cells derived from *Unc93b1*^*WT/WT*^, *Unc93b1*^*PKP/WT*^, and *Unc93b1*^*PKP/PKP*^ mice after stimulation with CpG-B (150 nM) for TLR9, R848 (10 ng/ml) and ssRNA40 (1 µg/ml) for TLR7, or LPS (10 ng/ml) for TLR4. (**e**) TNFα and IL-6 production, measured by ELISA and CBA respectively, of bone marrow-derived macrophages from the indicated mouse genotypes after stimulation for 8h with CpG-B (500 nM), LPS (50 ng/ml), or increasing concentrations of R848 (n=3). (**f**) Enhanced Myddosome assembly in macrophages from *Unc93b1*^*PKP/PKP*^ mice. Immunoprecipitation of Myd88 from bone marrow-derived macrophages from the indicated mice after stimulation with R848 (500 ng/ml), followed by immunoblot for IRAK2. Input levels of Myd88 and IRAK2 in whole cell lysates (WCL) are also shown. (**g**) TLR7 deficiency rescues disease in *Unc93b1*^*PKP/PKP*^ mice. Body weights of 6-week-old mice with indicated genotypes are shown in the left panel. Frequencies of CD44^high^ CD62L^neg^ of CD4^+^ T cells, and marginal zone (MZ) B cells (CD21^+^CD23^neg^IgM^+^CD1d^+^) in spleens are shown in the middle and right panels. Data points were pooled together from three independent experiments. All data are mean ± SD; *p < 0.05, **p < 0.01, ***p < 0.001 by unpaired Student’s t-test.

The phenotype of Unc93b1^PKP/PKP^ mice demonstrates that disruption of the Unc93b1/Syntenin-1 interaction decreases the activation threshold of TLR7, enabling responses to self RNA. To test this hypothesis, we examined the function of Unc93b1-dependent TLRs in cells from Unc93b1^PKP^-expressing mice. Bone marrow-derived dendritic cells (BMDCs) and macrophages (BMMs) from *Unc93b1*^*WT/PKP*^ and *Unc93b1*^*PKP/PKP*^ mice mounted stronger responses to TLR7 ligands compared to *Unc93b1*^*WT/WT*^ cells, while responses to TLR9 and TLR4 ligands were equivalent (Figs. 5d,e and S6b). Consistent with the model that Syntenin-1 alters the activation threshold of TLR7, enhanced responses to R848 were most evident at low ligand concentrations (Fig. 5e). In line with the enhanced cytokine production, macrophages from *Unc93b1*^*PKP/PKP*^ mice showed accelerated and stronger assembly of the Myddosome complex downstream of TLR7 activation (Fig. 5f). These enhanced TLR7 responses were not due to differences in Unc93b1 expression, as Unc93b1 protein levels were similar in BMMs from *Unc93b1*^*WT/WT*^, *Unc93b1*^*WT/PKP*^, and *Unc93b1*^*PKP/PKP*^ mice (Fig. S6c).

To test whether TLR7 function is required for the disease in *Unc93b1*^*PKP/PKP*^ mice we generated *Unc93b1*^*PKP/PKP*^*Tlr7*^*-/-*^ mice. Lack of TLR7 completely rescued disease as weight, numbers of activated T cells, numbers of MZ B cells, and frequencies of ANA were equivalent between *Unc93b1*^*PKP/PKP*^*Tlr7*^*-/-*^ and *Unc93b1*^*WT/WT*^ mice (Figs. 5c,g). These results indicate that Unc93b1 specifically limits TLR7-driven autoimmunity, presumably based on recognition of self-RNA, through recruitment of Syntenin-1.

These findings identify a new mechanism that specifically limits TLR7 signaling and consequently prevents responses against self-RNA. We propose a model whereby the C-terminal tail of Unc93b1 binds the negative regulator Syntenin-1, which is further recruited upon TLR7 activation and controls the threshold of TLR7 signaling (Fig. 6). Disrupting the binding site or recruitment of Syntenin-1 through mutations in Unc93b1 results in uncontrolled TLR7 responses that lead to a break in self-tolerance and autoimmunity. Syntenin-1 has been previously described as a versatile adaptor protein that regulates multiple signaling complexes in the cell. Our work suggests that Syntenin-1 can interfere with TLR7 signaling directly. This inhibition acts early in TLR7 signaling, affecting assembly of the Myddosome and subsequent downstream signaling. The precise mechanism by which Syntenin-1 inhibits Myddosome assembly remains unclear, but it seems likely that binding of Syntenin-1 to one or more signaling components prevents stable assembly of the multi-protein complex. Thus, Unc93b1 appears to serve as a regulatory scaffolding protein, bridging TLR7 with a negative regulator that prevents signal initiation.

**Fig. 6.**
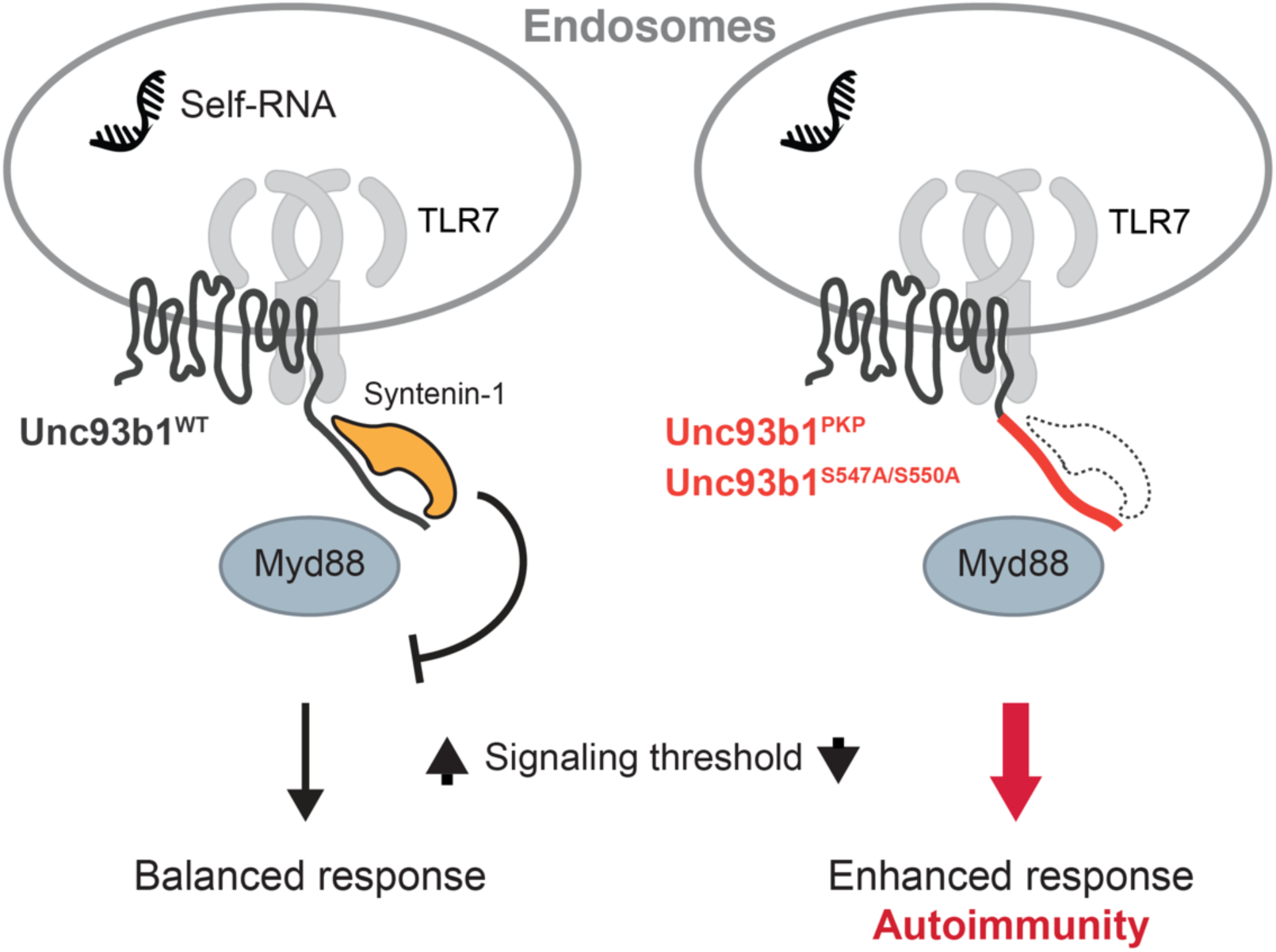
Model of Syntenin-1-mediated restriction of TLR7 signaling. The negative regulator Syntenin-1 binds to the C-terminal cytosolic tail of Unc93b1 to dampen TLR7 signaling and prevent responses to self RNA. Disruption of the Syntenin-1 binding site in Unc93b1 prevents Syntenin-1 recruitment and lowers the threshold for TLR7 activation, leading to TLR7-driven autoimmunity.

Previously described functions of Unc93b1 have been limited to control of TLR trafficking; our work clearly expands the scope of Unc93b1 function and suggests that this protein controls TLR function at multiple levels. One particularly interesting aspect of the mechanism we describe here is its selectivity for regulation of TLR7, especially considering the differential roles played by TLR7 and TLR9 in mouse models of SLE. The disease observed in *Unc93b1*^*PKP/PKP*^ mice was entirely TLR7 dependent, and there was no evidence of enhanced TLR9 signaling in these mice. In an accompanying manuscript^28^, we describe a mechanism that explains this selectivity: while Unc93b1 remains associated with TLR7 in endosomes, TLR9 is released and is therefore no longer subject to inhibition by Syntenin-1. Thus, our work provides the first mechanistic basis for differential regulation of TLR7 and TLR9, which may be relevant for human autoimmune diseases. We demonstrate that genetic variants in the C-terminal tail of human Unc93b1 can alter TLR7 responses, but even more intriguing is the possibility that Syntenin-1 recruitment to Unc93b1 can be dynamically controlled through phosphorylation or dephosphorylation of Ser547 and Ser550. Identifying the players involved in this regulation should reveal critical determinants that influence self versus non-self discrimination.

## Methods

### Antibodies and Reagents

The following antibodies were used for immunoblots and immunoprecipitations: anti-HA as purified antibody or matrix (3F10, Roche), anti-FLAG as purified antibody or matrix (M2, Sigma-Aldrich), anti-mLamp-1 (AF4320, R&D Systems), anti-Calnexin (ADI-SPA-860, Enzo Life Sciences), anti-Gapdh (GT239, GeneTex), anti-Myd88 (AF3109, R&D Systems), anti-IRAK2 (Cell Signaling), anti-Phospho-p38 (Cell Signaling), anti-p38 (Cell Signaling), anti-Phospho-SAPK/JNK (81E11, Cell Signaling), anti-SAPK/JNK (56G8, Cell Siganling), anti-Phospho-p44/42 (ERK1/2) (D13.14.4E, Cell Signaling), anti-p44/42 (ERK1/2) (137F5, Cell Siganling), anti-IκBα (Cell Signaling), anti-Syntenin-1 (2C12, Novusbio), anti-Unc93b1 (PA5-20510, Thermo Scientific), goat anti-mouse IgG-AlexaFluor680 (Invitrogen), goat anti-mouse IgG-AlexaFluor680 (Invitrogen), rabbit anti-goat IgG-AlexaFluor680 (Invitrogen), goat anti-mouse IRDye 800CW (Licor), donkey anti-rabbit IRDye 680RD (Licor), goat anti-rat IRDye 800CW (Licor). Antibodies for immunofluorescence were: rat anti-HA (3F10, Roche), rabbit anti-Lamp1 (ab24170, Abcam), goat anti-rat IgG-AlexaFluor488 (Jackson Immunoresearch), goat anti-rabbit IgG-AlexaFluor647 (Jackson Immunoresearch). Cells were mounted in Vectashield Hard Set Mounting Medium for Fluorescence (Vector Laboratories). For ELISA: anti-mouse TNFα purified (1F3F3D4, eBioscience), anti-mouse TNFα-biotin (XT3/XT22, eBioscience), Streptavidin-HRP (BD Pharmingen). Antibodies and reagents used for flow cytometry were: anti-TNFα (MP6-XT22, eBioscience), purified anti-CD16/32 Fc Block (2.4G2), CD3ε (145-2C11, BioLegend), CD4 (GK1.5, BioLegend), CD8 (53-6.7, BioLegend), CD44 (IM7, eBioscience), CD62L (MEL-14, eBioscience), CD69 (H1.2F3, eBioscience), CD1d (1B1, eBioscience), B220 (RA3-6B2, Invitrogen), CD19 (6D5, BioLegend), IgD (11-26c.2a, BioLegend), IgM (eB121-15F9, eBioscience), CD21 (eBio8D9, eBioscience), CD23 (B3B4, eBioscience), CD138 (281-2, BioLegend), CD11b (M1/70, BioLegend), Ly6G (1A8, TONBO biosciences), Ly6C (HK1.4, BioLegend), F4/80 (CI:A3-1, AbD serotec), MHCII (M5/114.15.2, eBioscience), CD86 (GL1, eBioscience), CD11c (N418, BioLegend). For ANA detection: anti-mouse IgG-AlexaFluor 488 (Jackson Immunoresearch), anti-mouse IgM-FITC (Invitrogen).

The antibody against phosphorylated Unc93b1 was generated by Invitrogen against synthesized phospho-peptide (YLEEDN(pS)DE(pS)DMEGEQ) using their “Rabbit, 90-Day immunization” protocol. Antibody in sera was enriched with immobilized phospho-peptide, followed by negative absorption with unphosphorylated peptide.

CpG-B (ODN1668: TCCATGACGTTCCTGATGCT, all phosphorothioate linkages) was synthesized by Integrated DNA Technologies. R848, PolyIC HMW, ssRNA40/LyoVec, and LPS were purchased from InvivoGen. Human IL-1b was from Invitrogen. NP-40 (Igepal CA-630) was from Sigma-Aldrich. Lipofectamine-LTX reagent (Invitrogen) and OptiMEM-I (Invitrogen) were used for transfection of plasmid DNA. ProMag 1 Series-COOH Surfactant free magnetic beads (#25029) for phagosome preparations were purchased from Polysciences. For luciferase assays: Renilla substrate: Coelenterazine native (Biotum), Firefly substrate: Luciferin (Biosynth), Passive Lysis Buffer, 5x (Promega).

### Animals

Mice were housed under specific-pathogen-free conditions at the University of California, Berkeley. All mouse experiments were performed in accordance with the guidelines of the Animal Care and Use Committee at UC Berkeley. Unless noted mice were analyzed at 5-8 weeks of age. C57BL/6J and TLR7^-/-^ mice (on the C57BL/6J background) were from the Jackson Laboratory. Unc93b1^PKP^ mice were generated using Cas9 genome editing. The guide RNA used was: TGCTGTGGCTTCGGAATGCGCGG. The single stranded oligo template contained 60bp homology arms on both sides and four phosphothioate linkages at the ends (one at the 5’ and three at the 3’ end of the oligo). Briefly, female C57BL/6J mice at 4 weeks of age were superovulated and mated overnight with C57BL/6J male mice (>8 weeks old). Zygotes were harvested from superovulated females and were placed in KSOM medium (Millipore) before use. CRISPR/Cas9 mixture was prepared in final concentration of *cas9* mRNA (100ng/ul), sgRNA (50ng/ul) and single stranded oligo (100ng/ul). The CRISPR/Cas9 mixture was microinjected into 80 zygotes using a micromanipulator (Narishige) and microscope (Nikon). After microinjection, 67embryos were transferred to three CD1 recipients via oviduct transfer. Offspring was genotyped by sequencing for the correct targeted allele and further bred to ensure germline transmission.

### Unc93b1 library design and plasmid constructs

The Unc93b1 mutagenesis library has been generated by Invitrogen. Briefly, the mouse Unc93b1 gene was optimized for the codon bias of *Mus musculus* and regions of very high (>80%) and very low (<30%) GC content have been avoided. The codon-optimized mouse Unc93b1 gene was c-terminally tagged with 3xFLAG (DYKDHDGDYKDHDIDYKDDDDK) and subjected to a triple-alanine scanning mutagenesis spanning sequences corresponding to tail and loop regions of the protein. The individual mutant constructs were cloned into a custom-made MSCV-based retroviral vector carrying an IRES-driven PuromycinR-T2A-mCherry double-selection. The library was provided as 204 individual plasmids.

For additional site-directed mutagenesis, AccuPrime Pfx DNA polymerase (Invitrogen) was used following the QuikChange II Site-directed Mutagenesis protocol from Agilent Technologies. The following MSCV-based retroviral vectors were used to express TLR7 and TLR9 in cell lines: MSCV2.2 (IRES-GFP), MSCV-Thy1.1 (IRES-Thy1.1), or MIGR2 (IRES-hCD2). TLR7 and TLR9 were fused to HA (YPYDVPDYA) at the C-terminal end. TLR7 sequence was synthesized after codon optimization by Invitrogen’s GeneArt Gene Synthesis service ^22^.

Syntenin-1 expression vector was from Addgene (89435). Traf6 expression vector was a kind gift from Jonathan Kagan (Boston Children’s Hospital).

### Cells and tissue culture conditions

HEK293T (from ATCC) and GP2-293 packaging cell lines (Clontech) were cultured in DMEM complete media supplemented with 10% (vol/vol) FCS, L-glutamine, penicillin-streptomycin, sodium pyruvate, and HEPES (pH 7.2) (Invitrogen). RAW264 macrophage cell lines (ATCC) were cultured in RPMI 1640 (same supplements as above). BMMs were differentiated for seven days in RPMI complete media (same supplements as above plus 0.00034% (vol/vol) beta-mercaptoethanol) and supplemented with M-CSF containing supernatant from 3T3-CSF cells, as previously described.

To generate HEK293T Unc93b1^-/-^ cells, guide RNAs were designed and synthesized as gBlocks as previously described^31^ and then were subcloned into pUC19 (guide RNA: CTCACCTACGGCGTCTACC). Humanized Cas9-2xNLS-GFP was a gift from the Doudna laboratory, University of California, Berkeley, CA. HEK293T cells were transfected using Lipofectamine LTX with equal amounts of the guide RNA plasmid and Cas9 plasmid. Seven days post transfection cells were plated in a limiting-dilution to obtain single cells. Correct targeting was verified by PCR analysis and loss of response to TLR9 and TLR7 stimulation in an NFkB luciferase assay. Unc93b1^-/-^ RAW macrophages were generated with the Cas9(D10A)-GFP nickase (guide RNAs: 1) GGCGCTTGCGGCGGTAGTAGCGG, 2) CGGAGTGGTCAAGAACGTGCTGG, 3) TTCGGAATGCGCGGCTGCCGCGG, 4) AGTCCGCGGCTACCGCTACCTGG). Macrophages were transfected with cas9(D10A) and all four guide RNAs using Lipofectamine LTX and Plus reagent and single cell-sorted on cas9-GFP two days later. Correct targeting was verified by loss of response to TLR7 stimulation and sequencing of the targeted region after TOPO cloning. Myd88 was knocked out in Unc93b1^-/-^ RAW macrophages stably expressing TLR7-HA and either Unc93b1^WT^ or Unc93b1^PKP^. Cas9 transfection and screening of cells was performed as before, except for using Cas9-2xNLS-GFP (guide RNA: GGTTCAAGAACAGCGATAGG).

### Retroviral transduction

Retroviral transduction of RAW macrophages was performed as previously described^22^. For macrophages expressing the Unc93b1 mutant library, transduced cells were selected with puromycin starting 48h after transduction and the efficiency of drug selection was verified by equal mCherry expression of target cells. When necessary, target cells were sorted on a Becton Dickinson Aria Fusion Sorter to match Unc93b1 expression levels using the bicistronic fluorescent reporter. For retroviral transduction of bone marrow derived macrophages, bone marrow was harvested and cultured in M-CSF-containing RPMI for two days. Progenitor cells were transduced with viral supernatant (produced as above) on two successive days by spinfection for 90 min at 32°C. 48h after the second transduction cells were put on Puromycin selection and cultured in M-CSF-containing RPMI media until harvested on day 8.

### Pulse-chase

Cells were seeded into 6 cm dishes the day before. After washing in PBS, cells were starved for 1 h in cysteine/methionine-free media (Corning) containing 10% dialyzed serum (dialyzed in PBS for two days using a 10 kD Snakeskin), then pulsed with 0.25 mCi/ml ^35^S-cysteine/methionine (EasyTag Express Protei Labeling Mix, Perkin-Elmer). After a 45-min pulse, cells were washed and cultured in 5 ml chase media containing 0.45 mg/ml L-cysteine and L-methionine or harvested as the zero time point. Time points were harvested as follows: cells were washed twice in 2 ml PBS, then scraped in PBS and cell pellets were subjected to HA immunoprecipitation.

### Cell fractionation by sucrose density-centrifugation

Cells in four confluent 15 cm dishes were washed in ice-cold PBS, scraped in 10 ml sucrose homogenization buffer (SHB: 250 µM sucrose, 3 mM imidazole pH 7.4) and pelleted by centrifugation. Cells were resuspended in 2 ml SHB plus protease inhibitor cocktail with EDTA (Roche) and 1mM PMSF and disrupted by 25 strokes in a steel dounce homogenizer. The disrupted cells were centrifuged for 10min at 1000g to remove nuclei. Supernatants were loaded onto continuous sucrose gradients (percent iodixanol: 0, 10, 20, 30) and ultracentrifuged in an SW41 rotor at 25800 rpm for 2 h (Optima L-90K Ultracentrifuge, Beckman Coulter). 22 fractions of 420 µl were collected from top to bottom. 100 µl of each fraction were denatured in SDS buffer for western blot analysis. For immunoprecipitations, three fractions corresponding to ER or endosomes were combined and lysed for 1h after addition of protease inhibitor cocktail and NP-40 to a final concentration of 1%. Coimmunoprecipitation with anti-HA matrix was performed as described below.

### Luciferase assays

Activation of NF-κB in HEK293T cells was performed as previously described^4^. Briefly, transfections were performed in OptiMEM-I (Invitrogen) with LTX transfection reagent (Invitrogen) according to manufacturer’s guidelines. Cells were stimulated with CpG-B (200 nM – 1 µM), R848 (100-200 ng/ml), or human IL-1b (20 ng/ml) after 24 h and lysed by passive lysis after an additional 12–16 h. Luciferase activity was measured on an LMaxII-384 luminometer (Molecular Devices).

### Immunoprecipitation, western blot, and dot blot

Cells were lysed in NP-40 buffer (50 mM Tris [pH 7.4], 150 mM NaCl, 0.5% NP-40, 5 mM EDTA, supplemented with 1mM PMSF, Roche complete protease inhibitor cocktail and PhosSTOP tablets). After incubation at 4°C for 1 h, lysates were cleared of insoluble material by centrifugation. For immunoprecipitations, lysates were incubated with anti-HA matrix or anti-FLAG matrix (both pre-blocked with 1% BSA-PBS) for at least 2 h, and washed four times in lysis buffer. Precipitated proteins were eluted in lysis buffer containing 200 ng/ml HA or 3xFLAG peptide, or denatured in SDS loading buffer at room temperature for 1 h. Proteins were separated by SDS-PAGE (Bio-Rad TGX precast gels) and transferred to Immobilon PVDF membranes (Millipore) in a Trans-Blot Turbo transfer system (Bio-Rad). Membranes were blocked with Odyssey blocking buffer, probed with the indicated antibodies and developed using the Licor Odyssey Blot Imager. For dot blot: diluted peptides were dropwise added to nitrocellulose blotting membranes (GE Healthware). Membranes were dried at room temperature, blocked and probed using the Licor Odyssey blot system.

Cell lysis and co-immunoprecipitations for Myddosome analyses were performed in the following buffer: 50 mM Tris-HCl pH 7.4, 150 mM NaCl, 10% glycerol, 1% NP-40 and supplemented with EDTA-free complete protease inhibitor cocktail (Roche), PhosSTOP (Roche) and 1 mM PMSF. Lysates were incubated overnight with anti-Myd88 antibody at 4°C, and then Protein G agarose (pre-blocked with 1% BSA-PBS) was added for additional 2 h. Beads were washed four times in lysis buffer, incubated in SDS loading buffer at room temperature for 1h, separated by SDS-PAGE, and probed with the indicated antibodies.

### Tissue harvest

Spleens and lymph nodes were digested with collagenase XI and DNase I for 30min and single cell suspensions were generated by mechanical disruption. Red blood cells were lysed in ACK Lysing Buffer (Gibco).

### Flow cytometry

Cells were seeded into non-treated tissue culture 24-well plates or round-bottom 96-well plates. The next day cells were stimulated with the indicated TLR ligands. To measure TNFα production, BrefeldinA (BD GolgiPlug, BD Biosciences) was added to cells 30 min after stimulation, and cells were collected after an additional 5.5 h. Dead cells were excluded using a fixable live/dead stain (Violet fluorescent reactive dye, Invitrogen). Cells were stained for intracellular TNFα with a Fixation & Permeabilization kit according to manufacturer’s instructions (eBioscience).

For flow cytometry on mouse cells, dead cells were excluded using a fixable live/dead stain (Aqua fluorescent reactive dye, Invitrogen) or DAPI and all stains were carried out in PBS containing 1% BSA (w/v) and 0.1% Azide (w/v) including anti-CD16/32 blocking antibody. Cells were stained for 20 min at 4°C with surface antibodies. Data were acquired on a LSRFortessa analyzer (BD Biosciences).

### Enzyme-linked immunosorbent assay (ELISA) and Cytometric bead array (CBA)

Cells were seeded into tissue culture-treated flat-bottom 96-well plates. The next day cells were stimulated with the indicated TLR ligands. For TNFα ELISAs, NUNC Maxisorp plates were coated with anti-TNFα at 1.5 µg/ml overnight at 4°C. Plates were then blocked with PBS + 1% BSA (w/v) at 37°C for 1 h before cell supernatants diluted in PBS + 1% BSA (w/v) were added and incubated at room temperature for 2 h. Secondary anti-TNFα-biotin was used at 1 µg/ml followed by Streptavidin-HRP. Plates were developed with 1 mg/mL OPD in Citrate Buffer (PBS with 0.05 M NaH_2_PO_4_ and 0.02 M Citric acid, pH 5.0) with HCl acid stop.

For CBA, cell supernatants were collected as above and analyzed using the Mouse Inflammation Kit (BD Biosciences) according to the manufacturer’s instructions.

### ANA staining

Mouse sera were diluted 1:80 in 1% BSA-PBS and applied to MBL Bion Hep-2 antigen substrate IFA test system for 1 h at room temperature. Slides were washed 3 times with PBS and incubated for 30 min with a mixture of fluorophore-conjugated secondary antibodies against anti-mouse IgG and IgM. Slides were washed 3 times and incubated with DAPI for 5 minutes. After rinsing once with PBS, slides were mounted in VectaShield Hard Set, and imaged on a Zeiss AxioZoom Z.1 slide scanner.

### Microscopy

Cells were plated onto coverslips and allowed to settle overnight. Coverslips were washed with PBS, fixed with 4% PFA-PBS for 15 min, and permeabilized with 0.5% saponin-PBS for 5 min. To quench PFA autofluorescence coverslips were treated with sodium borohydride/0.1% saponin-PBS for 10 min. After washing 3x with PBS, cells were blocked in 1% BSA/0.1% saponin-PBS for 1 h. Slides were stained in blocking buffer with anti-HA and anti-LAMP1 (see antibodies above), washed with PBS and incubated for 45 min with secondary antibodies. Cells were washed 3x in PBS and mounted in VectaShield Hard Set. Cells were imaged on a Zeiss Elyra PS.1 with a 100x/1.46 oil immersion objective in Immersol 518F / 30°C (Zeiss). Z-Sections were acquired, with three grid rotations at each Z-position. The resulting dataset was SIM processed and Channel Aligned using Zeiss default settings in Zen. The completed super-resolution Z-Series was visualized and analyzed using Fiji^32^. To compare the degree of colocalization of two proteins a single section from the middle of the Z-Series was selected and analyzed using a customized pipeline for object-based colocalization in Cell Profiler^33^. Briefly, primary objects (TLR7 vs Lamp1, or Unc93b1 vs Lamp1) were identified and related to each other to determine the degree of overlap between objects. Data are expressed as % of object 1 colocalized with object 2.

### Phagosome isolation and protein complex purification

Cells in a confluent 15cm dish were incubated with ∼10^8^ 1 µm magnetic beads (Polysciences) for 4 h. After rigorous washing in PBS, cells were scraped into 10 ml sucrose homogenization buffer (SHB: 250 µM sucrose, 3 mM imidazole, pH 7.4) and pelleted by centrifugation. Cells were resuspended in 2 ml SHB plus protease inhibitor cocktail with EDTA (Roche) and 1mM PMSF and disrupted by 25 strokes in a steel dounce homogenizer. The disrupted cells were gently rocked for 10 min on ice to free endosomes. Beads were collected with a magnet (Dynal) and washed 4x with SHB plus protease inhibitor. After the final wash, phagosome preparations were denatured in 2x SDS buffer at room temperature for 1 h and analyzed by western blot.

For protein complex purification, phagosome preparations were lysed in NP-40 buffer (50 mM Tris, pH 7.4, 150 mM NaCl, 0.5% NP-40, 5 mM EDTA, supplemented with 1 mM PMSF, complete protease inhibitor cocktail and PhosSTOP tablets (Roche) on ice for 1 h. Magnetic beads were removed by magnet and insoluble components were precipitated by 15,000 g spin for 20 min. Lysate was incubated with anti-FLAG matrix for 3 h, followed by four washes in lysis buffer. Proteins were eluted in NP-40 buffer containing 200 ng/ml 3xFLAG peptide, and were further applied to western blot, silver stain or Trypsin in-solution digest for mass spectrometry.

### Mass Spectrometry

Proteins were simultaneously extracted from a gel slice and digested with trypsin, and the resulting peptides were dried and resuspended in buffer A (5% acetonitrile/ 0.02% heptaflurobutyric acid (HBFA)). A nano LC column that consisted of 10 cm of Polaris c18 5 µm packing material (Varian) was packed in a 100 µm inner diameter glass capillary with an emitter tip. After sample loading and washed extensively with buffer A, the column was then directly coupled to an electrospray ionization source mounted on a Thermo-Fisher LTQ XL linear ion trap mass spectrometer. An Agilent 1200 HPLC equipped with a split line so as to deliver a flow rate of 300 nl/min was used for chromatography. Peptides were eluted using a 90 min. gradient from buffer A to 60% Buffer B (80% acetonitrile/ 0.02% HBFA).

Protein identification and quantification were done with IntegratedProteomics Pipeline (IP2, Integrated Proteomics Applications, Inc. San Diego, CA) using ProLuCID/Sequest, DTASelect2 and Census. Tandem mass spectra were extracted from raw files using RawExtractor and were searched against the mouse protein database (obtained from UNIPROT) plus sequences of common contaminants, concatenated to a decoy database in which the sequence for each entry in the original database was reversed. LTQ data was searched with 3000.0 milli-amu precursor tolerance and the fragment ions were restricted to a 600.0 ppm tolerance. All searches were parallelized and searched on the VJC proteomics cluster. Search space included all fully tryptic peptide candidates with no missed cleavage restrictions. Carbamidomethylation (+57.02146) of cysteine was considered a static modification. We required 1 peptide per protein and both trypitic termini for each peptide identification. The ProLuCID search results were assembled and filtered using the DTASelect program with a peptide false discovery rate (FDR) of 0.001 for single peptides and a peptide FDR of 0.005 for additional peptide s for the same protein. Under such filtering conditions, the estimated false discovery rate was zero for the datasets used.

### Quantification and Statistical Analysis

Statistical parameters, including the exact value of n and statistical significance, are reported in the Figures and Figure Legends, whereby n refers to the number of repeats within the same experiment. Representative images have been repeated at least three times, unless otherwise stated in the figure legends. Data is judged to be statistically significant when p < 0.05 by Student’s t-test. To compare the means of several independent groups, a one-way ANOVA followed by a Tukey’s posttest was used. In figures, asterisks denote statistical significance (*, p < 0.05; **, p < 0.01; ***, p < 0.001). Statistical analysis was performed in GraphPad PRISM 7 (Graph Pad *Software* Inc.).

## Acknowledgements

We thank Russell Vance and members of the Barton and Vance Labs for helpful discussions and critical reading of the manuscript. We thank Angus Yiu-fai Lee and the Gene Targeting Facilty of the Cancer Research Center at UC Berkeley for generating the *Unc93b1*^*PKP/PKP*^ knock-in mice. We thank Lori Kohlstaedt and the Vincent J. Coates Proteomics/Mass Spectrometry Laboratory at UC Berkeley for identification of Unc93b1 interacting proteins. We thank Hector Nolla and Alma Valeros for assistance with cell sorting at the Flow Cytometry Facility of the Cancer Research Laboratory at UC Berkeley. We thank Steven Ruzin and Denise Schichnes for assistance with microscopy on the Zeiss Elyra PS.1 at the Biological Imaging Center at UC Berkeley. This work was supported by the NIH (AI072429, AI105184 and AI063302 to G.M.B.) and by the Lupus Research Institute (Distinguished Innovator Award to G.M.B.). O.M. was supported by an Erwin Schrödinger (J 3415-B22) and CRI Irvington postdoctoral fellowship. B.L. was supported by the UC Berkeley Tang Distinguished Scholars Program. Research reported in this publication was supported in part by the NIH S10 program under award number 1S10OD018136-01 and by the NIH S10 Instrumentation Grant S10RR025622.

## Author Contributions

O.M., B.L., and G.M.B designed experiments. O.M. and B.L. performed experiments and analyzed the data. N.K. designed experiments and analyzed data related to mass spectrometry. G.M.B. wrote the manuscript. O.M and B.L. revised and edited the manuscript.

## Supplementary Materials for

**Fig. S1:**
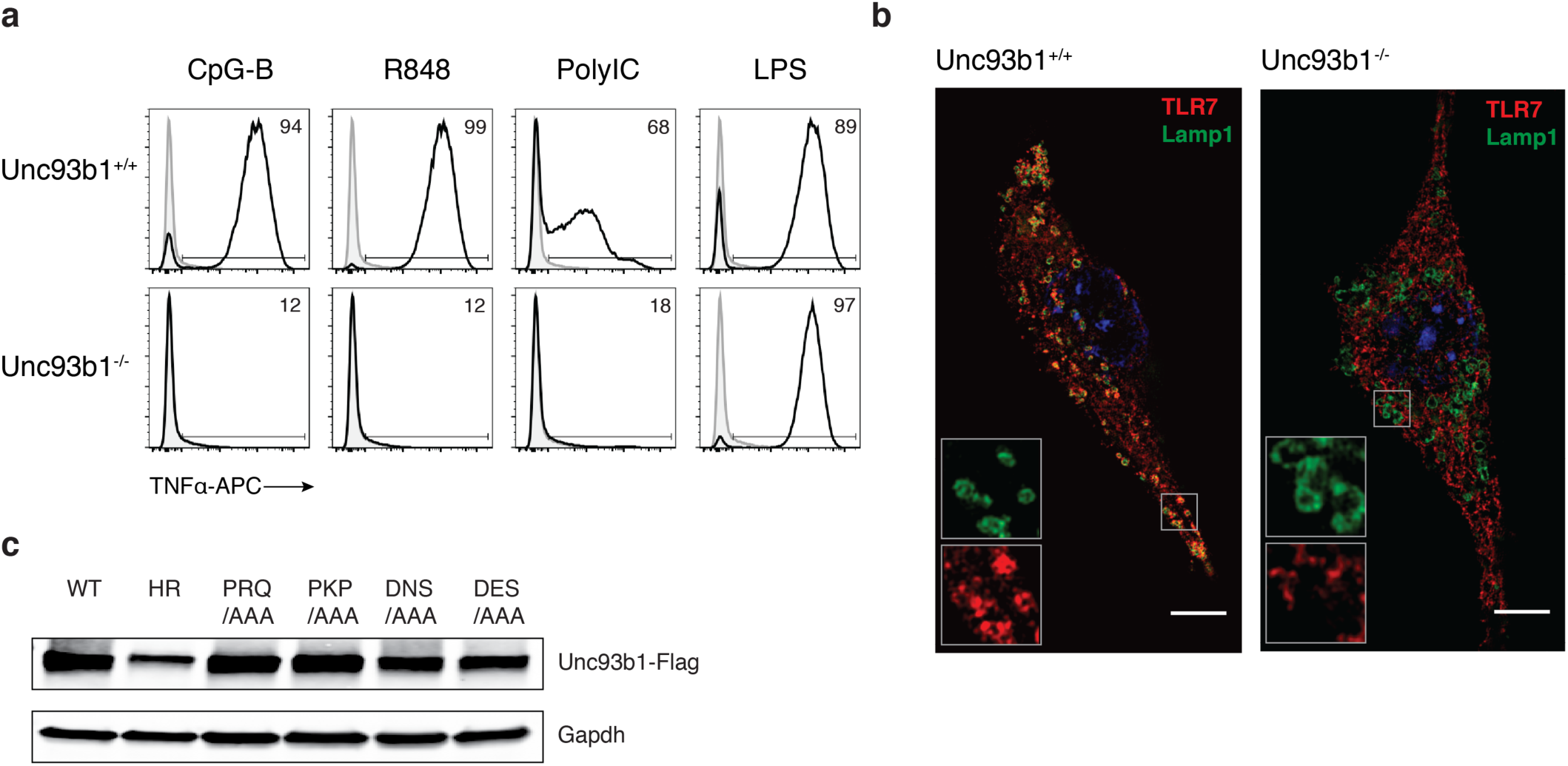
Generation and validation of Unc93b1-deficient RAW macrophages. (**a**) Unc93b1-deficient RAW macrophages are unresponsive to nucleic acid ligands. Representative flow cytometry analysis showing percent TNFα positive cells, measured by intracellular cytokine staining, of WT and Unc93b1-deficient RAW macrophage line, generated by CRISPR/Cas9, after stimulation with CpG-B (1 µM) for TLR9, R848 (400 ng/ml) for TLR7, PolyIC (20 µg/ml) for TLR3, or LPS (10 ng/ml) for TLR4. (**b**) TLR7 does not traffic to endosomes in Unc93b1-deficient RAW macrophages. Colocalization of TLR7 and Lamp1 in RAW macrophages expressing the indicated Unc93b1 alleles using superresolution structured illumination microscopy. Representative images are shown with TLR7 (red) and Lamp1 (green). (**c**) Unc93b1 expression levels, as measured by FLAG immunoblot, of Unc93b1-deficient RAW macrophages retrovirally transduced to express the indicated Unc93b1 alleles. These same cell lines are used for experiments shown in Fig. 1a. The data are representative of at least three independent experiments, unless otherwise noted.

**Fig. S2:**
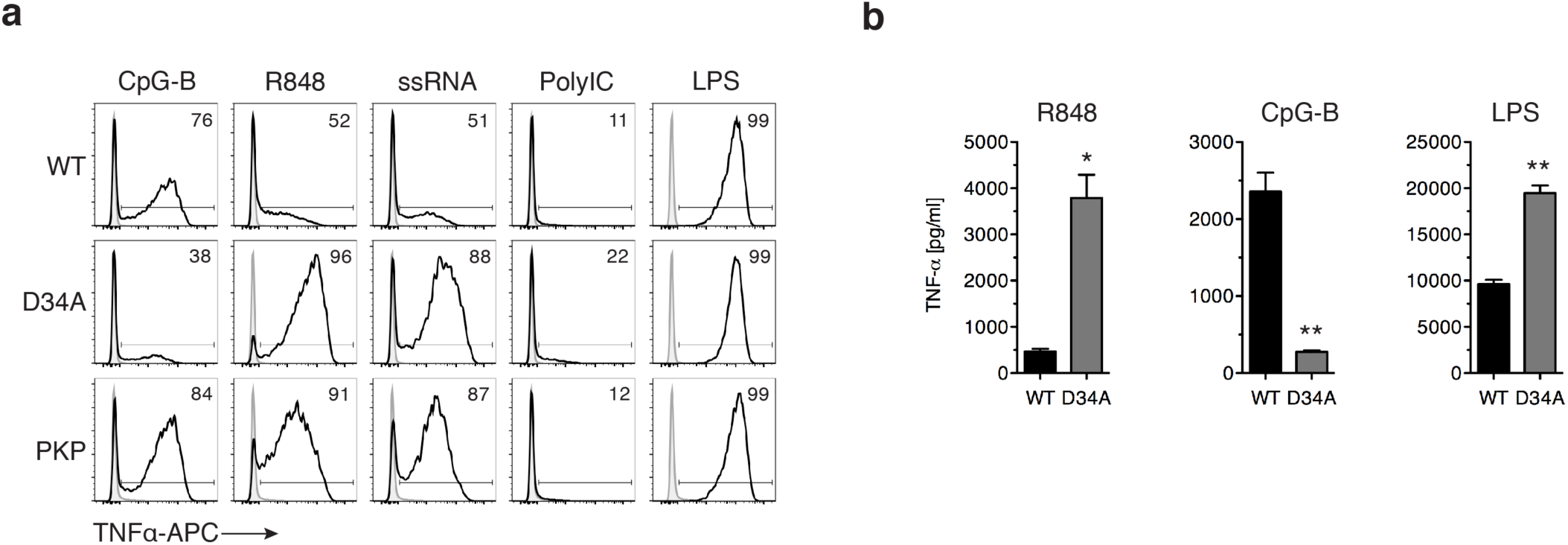
Unc93b1^PKP^ does not alter TLR9 responses, unlike Unc93b1^D^34^A^. (**a**) Representative flow cytometry analysis showing percent TNFα positive cells, measured by intracellular cytokine staining, of indicated RAW macrophage lines after stimulation with CpG-B (25 nM) for TLR9, R848 (10 ng/ml) and ssRNA40 (1 µg/ml) for TLR7, PolyIC (20 µg/ml) for TLR3, or LPS (10 ng/ml) for TLR4. Representative of three independent experiments. (**b**) TNFα production, measured by ELISA, from the indicated RAW macrophage lines after stimulation for 8h with R848 (10 ng/ml), CpG-B (25 nM), or LPS (50 ng/ml) (n=3, representative of two independent experiments). All data are mean ± SD; *p < 0.05, **p < 0.01, ***p < 0.001 by unpaired Student’s t-test.

**Fig. S3:**
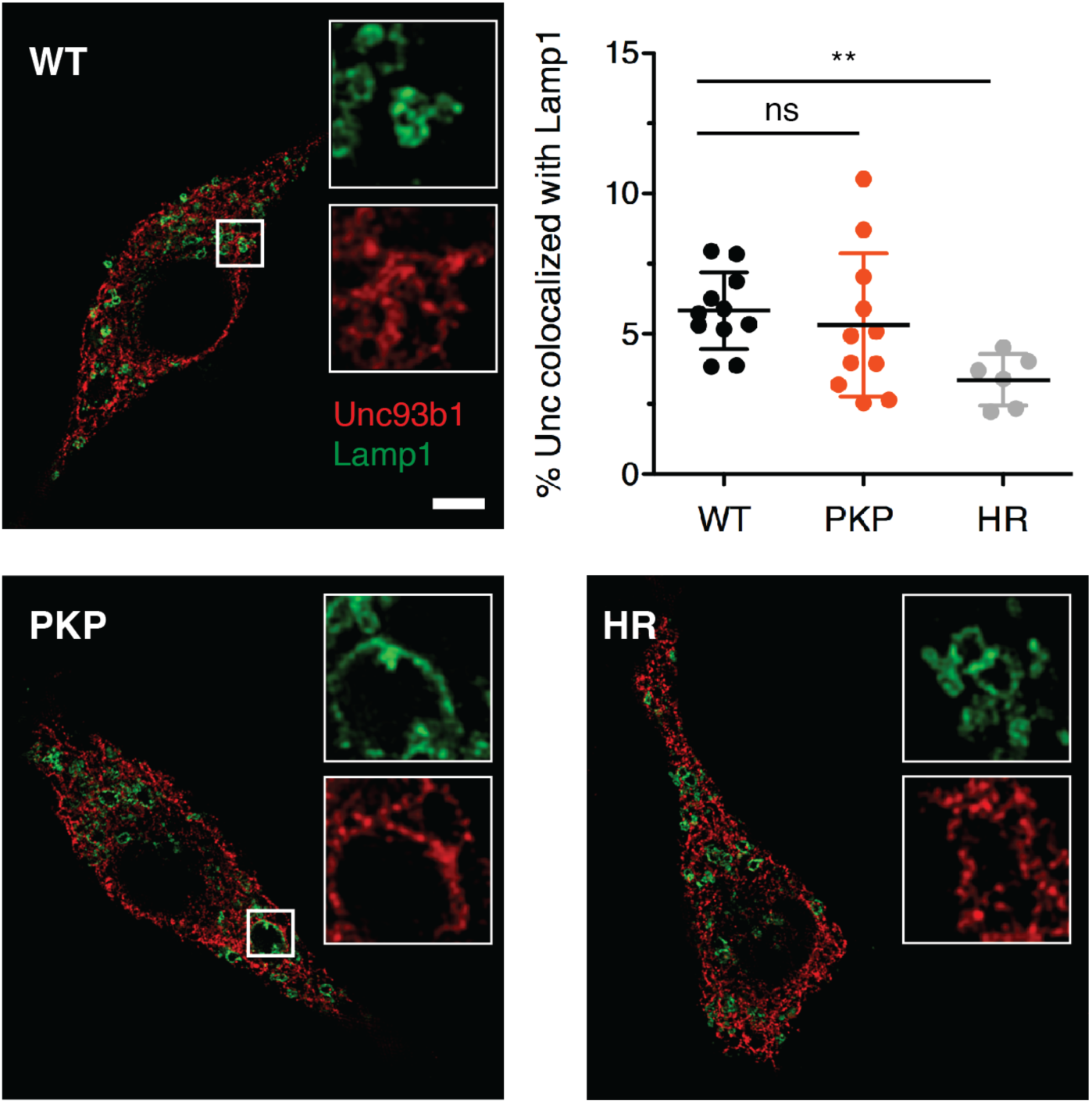
The sub-cellular localization of Unc93b1^PKP^ is not altered relative to Unc93b1^WT^. Colocalization of Unc93b1 (red) and Lamp1 (green) was measured using superresolution structured illumination microscopy in Unc93b1-deficient RAW macrophages complemented with Unc93b1^WT^, Unc93b1^PKP^, or Unc93b1^H412R^. A representative cell is shown for each Unc93b1 allele. Boxed areas are magnified. The plot shows quantification of the percentage of total Unc93b1 within Lamp1^+^ endosomes with each dot representing an individual cell (imaged in a single experiment). Scale bars: 10µm. Data is presented as mean ± SD; **p < 0.01 by unpaired Student’s t-test.

**Fig. S4:**
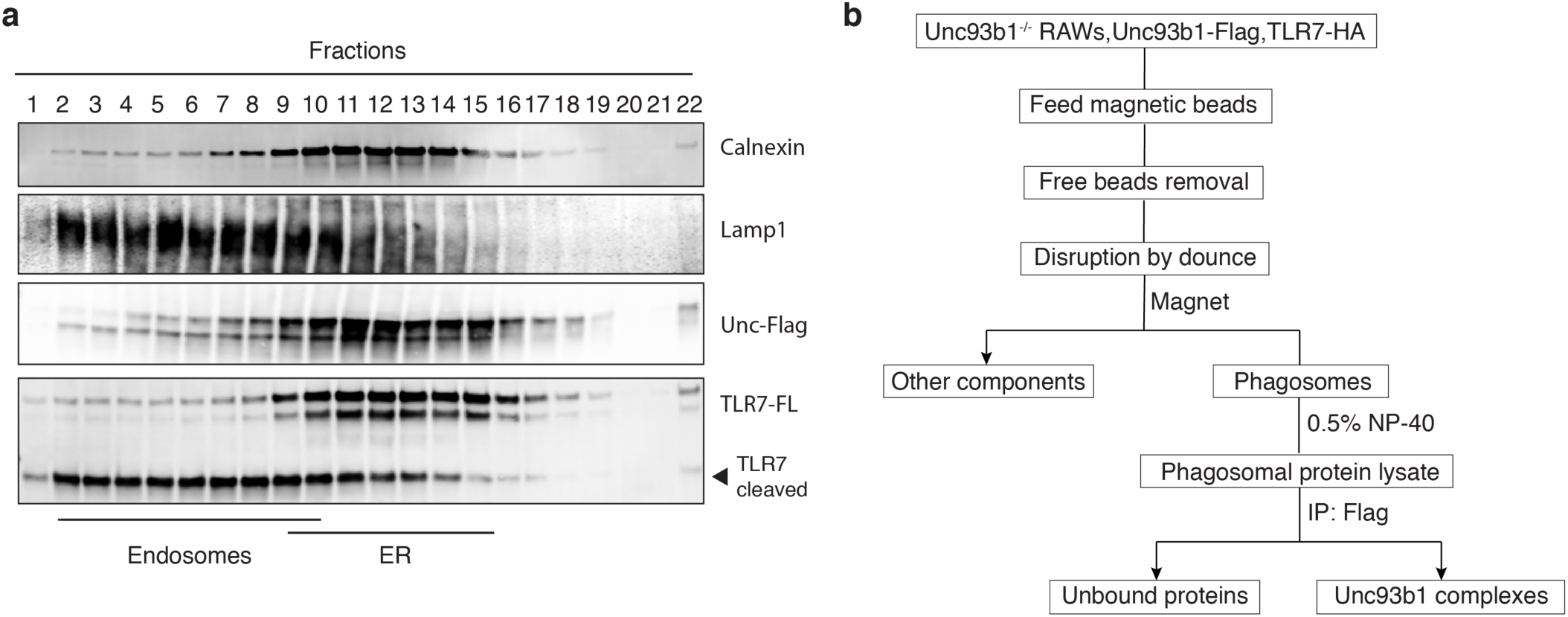
Mass spectrometry analysis of Unc93b1 complexes. (**a**) A small fraction of Unc93b1 resides in endosomes compared to the endoplasmic reticulum (ER). Sub-cellular fractionation of TLR7-HA, Unc93b1-FLAG expressing RAW macrophages was performed by density-gradient centrifugation. The distribution of Calnexin (ER), Lamp1 (late endosomes and lysosomes), Unc93b1, and TLR7 across fractions was measured by immunoblot. Representative of three independent experiments. (**b**) Workflow for isolation of phagosomes from RAW macrophages and purification of Unc93b1-FLAG complexes from phagosome lysates.

**Fig. S5:**
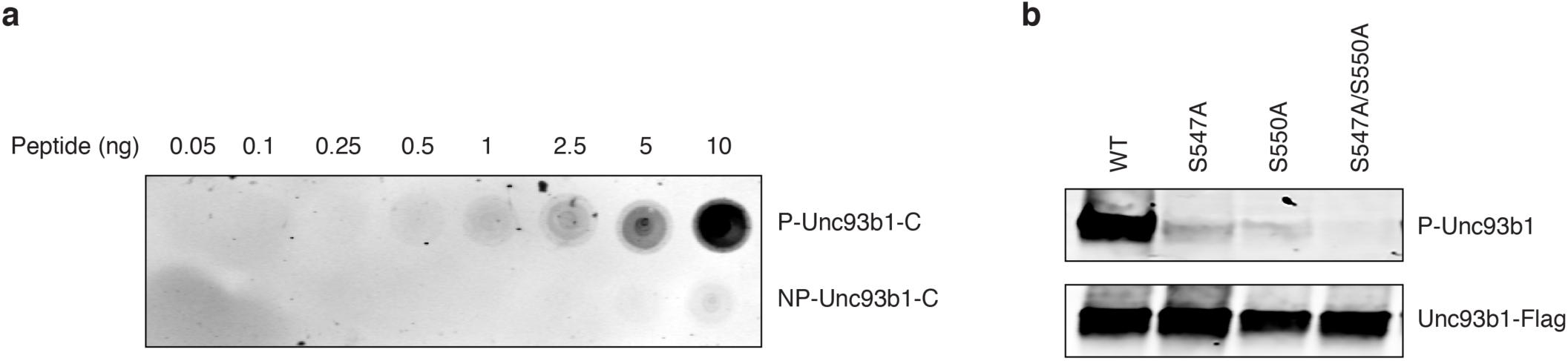
Validation of the anti-phospho-Unc93b1 polyclonal Unc93b1 antibody. **(a)** Immunoblots demonstrating the specificity of the phosphospecific antibodies generated against Ser547 and Ser550 in the Unc93b1 C-tail. Varying quantities of synthesized peptides corresponding to the Unc93b1 C-terminal regulatory region with (P-Unc93b1-C) and without (NP-Unc93b1-C) phosphorylated Ser547 and Ser550 were blotted to membrane and probed with rabbit phospho-specific, affinity-purified polyclonal IgG. Representative of two independent experiments. (**b**) The phospho-specific polyclonal antibody detects both phosphorylated Ser547 and Ser550. Unc93b1 was isolated from Unc93b1-deficient RAW macrophages expressing Unc93b1^S^547^A^, Unc93b1^S^550^A^, or Unc93b1^S^547^A/S^550^A^ by FLAG immunoprecipitation followed by immunoblot with the phospho-specific polyclonal antibody. Representative of at least three independent experiments.

**Fig. S6:**
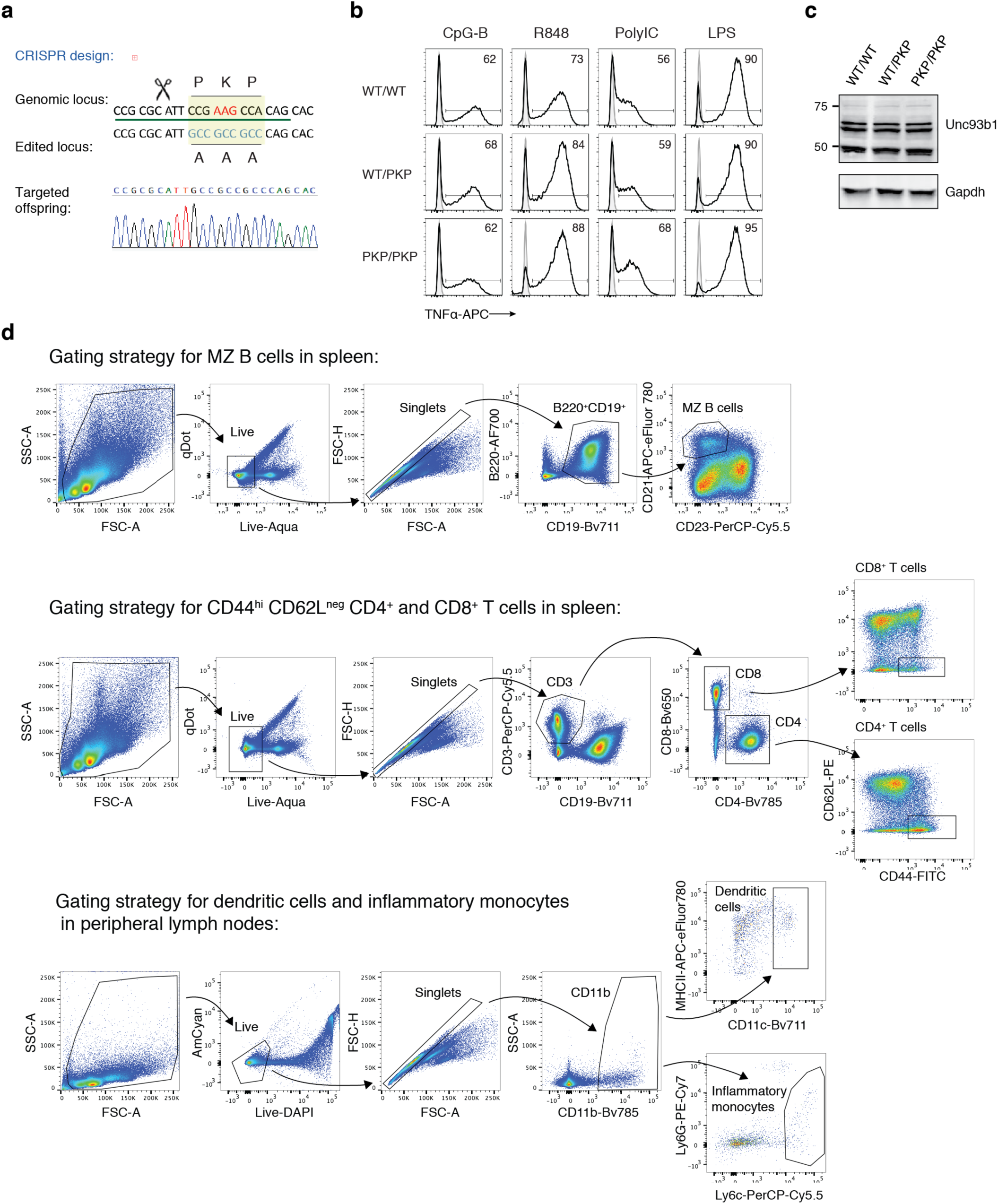
Unc93b1^PKP^ knock-in mice develop systemic inflammation. (**a**) CRISPR/Cas-9 strategy to generate Unc93b1^PKP^ knock-in mice. (**b**) Representative flow cytometry analysis showing percent TNFα positive cells, measured by intracellular cytokine staining, of bone marrow-derived macrophages derived from *Unc93b1*^*WT/WT*^, *Unc93b1*^*PKP/WT*^, and *Unc93b1*^*PKP/PKP*^ mice after stimulation with CpG-B (150 nM) for TLR9, R848 (10 ng/ml) for TLR7, PolyIC (10 µg/ml) for TLR3, or LPS (10 ng/ml) for TLR4. **(c)** Unc93b1 protein levels in bone marrow-derived macrophages from indicated mouse genotypes, measured by immunoblot with polyclonal antibodies against endogenous Unc93b1. **(d)** Representative gating strategies for marginal zone (MZ) B cells, activated T cells, dendritic cells and inflammatory monocytes are shown. These strategies were used for the data presented in Figure 5.

